# Cell-associated viral community composition and its functional potential in a dimictic lake on the Canadian Shield

**DOI:** 10.1101/2024.12.22.629985

**Authors:** Cody C. Collis, Ellen S. Cameron, Kirsten M. Müller, Monica B. Emelko, Nicolas Tromas, Jozef I. Nissimov

## Abstract

The Turkey Lake Watershed (TLW), located in Northern Ontario, forms a cascading lake system that ultimately drains into Lake Superior. Historically, the TLW has served as a key research site for studies on acid rain and climate change. Despite its long-standing ecological importance, investigations into its microbial communities remain limited, and to date, no studies have explicitly examined the viral component of this watershed. A major obstacle to such research is the challenge of obtaining sufficient viral biomass from remote locations like the TLW for downstream sequencing and analysis. To address this, we focused on viral signatures associated with microbial cells, which require less biomass at the time of collection. Using this approach, we conducted a seasonal metagenomic survey of Big Turkey Lake (BTL)—one of the primary lakes within the TLW—from summer 2018 to winter 2020. Our study characterized the diversity and composition of cell-associated viral communities, predicted potential host relationships, and identified auxiliary metabolic genes (AMGs) to better understand the functional roles of these viruses. The results revealed a dynamic viral community that varied across seasons. Most viral contigs were classified within the class *Caudoviricetes*, while members of the *Phycodnaviridae* emerged as important non-bacteriophage contributors to the viral assemblage. We also reconstructed six draft viral metagenome-assembled genomes (vMAGs), ranging in size from 11 to 45 kbp, all identified as *Caudoviricetes*, with two being potential temperate phages, and a third containing ribosomal genes. Notably, AMGs associated with lipid metabolism were detected exclusively in a winter sample, suggesting a potential season- or niche-specific role for these genes in BTL, while AMGs related to phosphorus acquisition were present in all samples. Together, these findings provide the first insight into the viral ecology of the TLW and establish a foundation for future studies investigating the roles of viruses in shaping the microbial dynamics in this important habitat.

## 1 Introduction

Viruses are integral components of microbial communities in aquatic environments, where they play crucial roles in biogeochemical cycling, regulation of primary production, and bacterial mortality (Suttle, 2007; Mojica *et al*., 2016; Mojica & Brussaard, 2014). By infecting and lysing host cells, viruses influence nutrient turnover and energy flow. They also shape bacterial and microalgal activity through the expression of auxiliary metabolic genes (AMGs)—genes acquired via evolutionary processes such as horizontal gene transfer during infection (Breitbart *et al*., 2007). Although these genes typically do not play a direct role in virus replication, they can enhance host metabolism, affecting key cellular functions. In aquatic systems, AMGs are implicated in processes such as photosynthesis, carbon metabolism, cofactor biosynthesis, and nutrient cycling (Hurwitz & U’ren, 2016; Crummett *et al*., 2016; Meza-Padilla *et al*., 2024). Notably, approximately 40% of psbA (photosystem II) expression in marine systems is virus-derived (Sieradzki *et al*., 2019), underscoring the ecological significance of viruses and their AMGs.

While the majority of research on aquatic viruses has focused on marine ecosystems, recent advances have revealed a remarkable diversity of viral and microbial communities in freshwater environments (Thompson et al., 2017; *Zhou et al*., 2025). Seasonal patterns in viral abundance and community composition are largely governed by host availability, which is influenced by bottom-up factors such as nutrient supply and top-down factors such as predation (Zhou *et al*., 2025). In typical freshwater lakes, seasonal phytoplankton succession—for example, shifts between diatoms and cyanobacteria driven by changes in temperature, light, and nutrient conditions—can strongly influence the composition of viral populations (Wentzky *et al*., 2020; Wei *et al*., 2020).

Despite these advances, major gaps persist in our understanding of viral infection dynamics and metabolic capabilities in most aquatic environments (Sime-Ngando, 2014; Bonetti *et al*., 2019; Pradeep Ram & Sime-Ngando, 2023). One area of uncertainty is the role of viruses during harmful algal blooms (HABs). Some studies suggest that viruses contribute to bloom termination (Grasso et al., 2022), while others indicate that viral infections can exacerbate HAB effects—viral lysis may release intracellular toxins into the water column, facilitating their persistence in dissolved form (Steffen et al., 2017; McKindles et al., 2020; Lee et al., 2023). The function of AMGs during HAB events also remains poorly understood. For instance, a cyanophage AMG involved in host phycobilisome degradation may help sustain infection under high light and nutrient-limited conditions (Meza-Padilla et al., 2024), but the prevalence of this and similar genes in freshwater viral communities is still unclear.

Although oceanic systems have provided a strong foundation for understanding viral ecology, knowledge of viral impacts in freshwater systems has lagged behind. Differences in scale and physical structure mean that findings from marine studies cannot always be extrapolated to lakes. Freshwater systems are generally shallower and enclosed, making them more sensitive to local factors such as eutrophication and climate change (Adrian et al., 2009; Williamson et al., 2008). Understanding viral roles in these ecosystems is thus essential for predicting how microbial communities respond to environmental stressors.

A crucial first step toward this understanding is characterizing the composition and relative abundance of viruses in freshwater environments, including those associated with host cells. However, sampling viruses in these settings poses practical challenges, particularly in remote habitats such as inland lakes of Northern Ontario. Unlike oceanic expeditions where onboard facilities allow collection of large water volumes, inland lake sampling is often limited by equipment and accessibility, making traditional virus concentration methods such as TFF of large volumes (Wommack *et al*., 2010) or chemical flocculation (John et al., 2011), impractical. An alternative is to focus on the cell-associated viral fraction, which includes viruses attached to or infecting host cells. This approach simplifies field collection, requiring only filtration of host cells rather than extensive water concentration procedures.

Here, we used shotgun metagenomics to characterize seasonal variation in the relative abundance of actively infecting or cell-associated viruses in Big Turkey Lake, a dimictic lake which is part of the Turkey Lake Watershed (TLW) in Northern Ontario, Canada. We also investigated potential viral hosts and identified AMGs that may mediate host metabolic processes. To our knowledge, this represents the first study of viral communities in the TLW, establishing a foundation for future research into the ecological roles and functional potential of freshwater viruses in this watershed.

## 2 Materials & Methods

### 2.1 Study site and sampling

This study was conducted using samples obtained between 2018 and 2020 (Table S1) from Big Turkey Lake (BTL; 47 02’ 54.7” North, 84 25’ 19.3” West) within the Turkey Lake Watershed (TLW; Figure 1-A), located approximately 50 kilometers north of Sault Ste. Marie, Ontario. The watershed, which consists of several holomictic lakes (i.e., lakes which experience complete water column mixing in spring and autumn), represents a relatively undisturbed habitat by human activity, and has been the prior focus of studies on the effects of acid rain and other aspects related to climate change (Jeffries & Foster, 2001). BTL is the largest lake within the TLW and is classified as dimictic (e.g., its water fully mixes top-to-bottom usually twice yearly) and oligotrophic (Jeffries *et al*., 1988). Briefly, 1 L water samples (Table S1) were vacuum filtered through 47 mm (1.2 µm) Whatman GF/C filters to capture microbial biomass, then kept at -20°C until downstream processing (Cameron, 2021).

**Figure 1.**
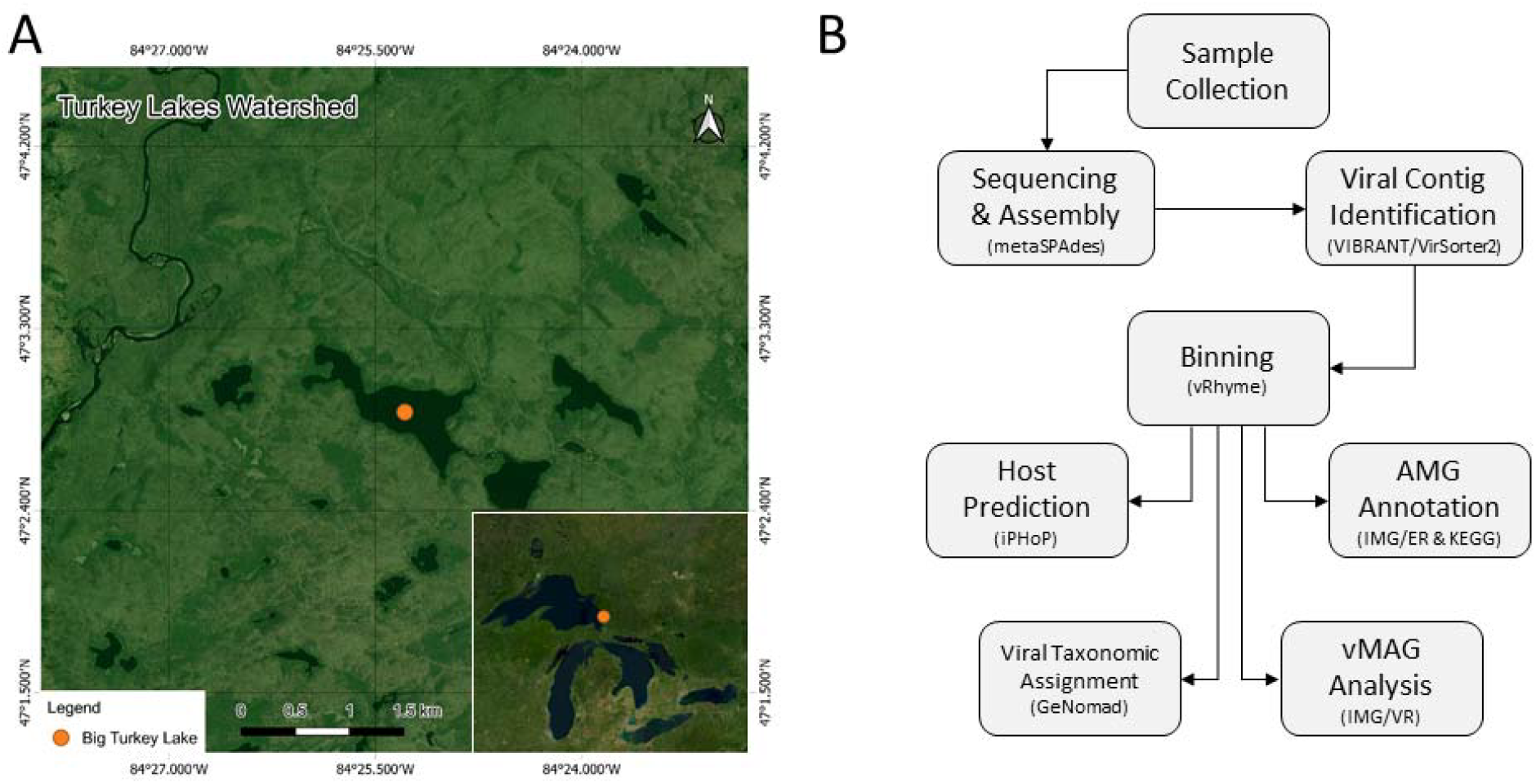
The sampling location of this study was Big Turkey Lake (BTL), within the Turkey Lake Watershed (TLW), which is adjacent to Lake Superior, Canada (**A**). The study consists of biomass samples collected during July 2018 – January 2020 for shotgun metagenomics. Extracted nucleic acids were subject to sequencing and a bioinformatics workflow (**B**) for assessing the microbial cell-associated viral communities.

### 2.2 Sequencing, assembly, and analysis

For samples taken in 2018, DNA extraction was performed using the Qiagen DNeasy PowerSoil kit, with shotgun sequencing being done by Metagenom Bio. Inc (Waterloo, Ontario) with Illumina MiSeq 250x2 paired-end sequencing. For samples taken in 2019 and 2020, DNA extraction and shotgun sequencing was performed by the University of Toronto’s Centre for the Analysis Genome Evolution and Function using the Qiagen DNeasy PowerSoil kit, and Illumina MiSeq 250x2 paired-end sequencing, respectively. Raw reads (Table S2) were subsequently put through the Metagenome-atlas pipeline (ver. 2.9.0) for both quality control and assembly, with MetaSPAdes as the assembler (Kieser *et al*., 2020; Nurk *et al*., 2017). Bioinformatics analysis was as described below and in Figure 1-B.

### 2.3 Virus identification and taxonomic classification

To identify the viral contigs, a combination of VIBRANT and VirSorter2 was used (Kieft *et al*., 2020; Guo *et al*., 2021) from within the WhatThePhage pipeline (ver. 1.1.0) (Marquet *et al*., 2022). Viral contigs were then binned using vRhyme (ver. 1.1.0) (Kieft *et al*., 2021), with the resulting viral metagenome-assembled genomes (vMAGs) assessed for completeness using CheckV (ver. 0.9.0) (Nayfach *et al*, 2021). Those vMAGs that had a completeness score of >90% from CheckV were considered as high-quality MAGs (Nayfach *et al*., 2021) and used for further analysis. Taxonomic evaluation of the viral contigs was performed using the annotate function of geNomad (ver. 1.11.1) (Camago *et al*., 2023) using default parameters, while relative abundance was calculated based on the number of viral contigs in each sample. Protein annotation for viral contigs was performed using JGI’s Integrated Microbial Genomes with Microbiome Samples – Expert Review (IMG/ER) resources (Chen *et al*., 2021). Coverage of the viral contigs was performed using CoverM (ver. 0.6.1) with the contig function, using the --*rpkm, --min-covered-fraction 30*, and *--proper-pairs-only* options (Aroney *et al*., 2024), and subsequent rarefaction analysis (Figure S1) was performed using the *ggrarecurve* function from the MicrobiotaProcess R package (ver.1.15.0) (Xu *et al*., 2023). For vMAGs that were scored as ‘complete’ by CheckV, visualizations were created using *gbdraw* (Kawato, 2026).

### 2.4 Host predictions and auxiliary metabolic gene analysis

To determine auxiliary metabolic genes (AMGs) found within the BTL cell-associated virus community, IMG/ER annotations were compared with the KEGG Orthology (KO) database to determine the specific functional potential of each gene (Kanehisa *et al*., 2016). Annotations were defined as an auxiliary metabolic gene (AMG) if they were not directly involved with viral replication, including DNA synthesis and transcription. Those that were defined as an AMG had their flanking regions examined to determine the presence of viral hallmark genes. Host prediction for all viral contigs was performed using the iPHoP pipeline using the default host database (ver. 1.3.1) (Roux, 2023), which has been designed to predict bacterial and archaeal hosts of phages based on their genomic sequences within viral contigs.

## 3 Results and Discussion

### 3.1 Cell-associated viral composition and abundance

In our study, hosts were collected on 1.2 µm filters, a pore size too large to capture most free-floating viruses. The focus is therefore on viruses that were associated with cells (i.e., viruses that were either actively infecting host cells or were attached to their exterior), which resulted in a total of 2.8-4.54 M raw reads (Table S2) with sufficient coverage for analysis (Figure S1). Across all samples, approximately 65% of total viral contigs were able to be taxonomically classified through geNomad analysis. The remaining viral contigs (approx. 35%) were unable to be classified and represent potentially novel viral sequences.

Within the identified viral contigs in BTL, there were two major classes of viruses represented (Figure 2-A): *Caudoviricetes*, otherwise known as bacteriophages, and *Megaviricetes*, large double-stranded DNA viruses that are part of the nucleocytoplasmic large DNA viruses (NCLDVs), capable of infecting a diverse range of eukaryotic organisms (Koonin & Yutin, 2010). In BTL, *Caudoviricetes* displayed the highest relative abundance (RA) of all viruses (57.5% of total viral contigs) and were present in all samples. *Megaviricetes* were also present in all samples, but with a comparatively lower RA (7.3% of total viral contigs). This is in accordance with other studies that have found that bacteriophages are generally more abundant than their NCLDV counterparts in lake environments. For example, in a study of Lake Ontario and Lake Erie, viruses belonging to the previously assigned *Myoviridae* family of bacteriophages comprised ∼80% of the viral sequences detected, whereas phycodnaviruses, members of *Megaviricetes*, made up a much smaller portion of the viral community (Mohiuddin and Schellhorn, 2015). Nevertheless, *Megaviricetes*, while less abundant overall, play a significant role in modulating phytoplankton dynamics and influence community structure and succession in most aquatic environments (Gimenes *et al*., 2011). Importantly, the abundance and diversity of bacteriophages and *Megaviricetes* can be influenced by factors such as the trophic state of a lake, its depth, and seasonal variations such as temperature. In our study, the RA of identified *Megaviricetes* was highest during the summer months of June and August of 2019 (24.4% and 16.7%, respectively). The reverse was seen in May 2019 and January 2020, where the RA of identified *Megaviricetes* were 1.5% and 1.8% respectively. *Caudoviricetes* display the reverse trends in their RA, with summer abundances being reduced to 25% in August 2019 and increasing to approximately 70% in both May 2019 and January 2020. The summer months at BTL typically have temperatures higher than 20°C whereas winter is typically below freezing with ice cover (Table S1). Hence, given that microbial host abundance and composition are known to be impacted by temperature in most aquatic systems, we conclude that temperature naturally also plays a central role in structuring the cell-associated viral abundance and its composition in BTL.

**Figure 2.**
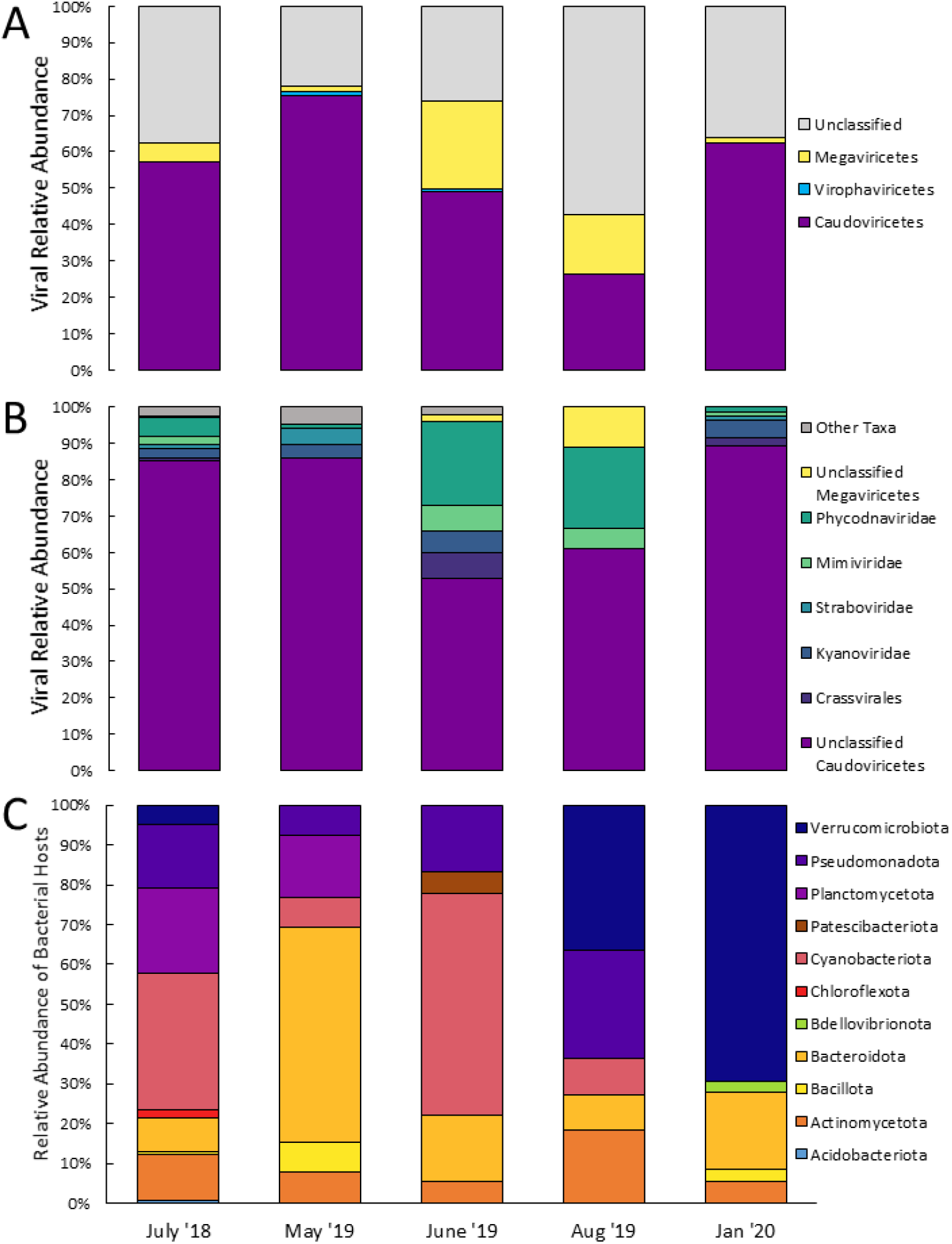
Taxonomy of cell-associated viruses and their respective hosts within Big Turkey Lake. **(A)** Relative abundance of cell associated virus community by class. **(B)** Relative abundance of cell-associated virus community by family, omitting unclassified sequences. **(C)** Host prediction analysis using IPHoP for cell associated classified and unclassified viruses across seasons.

Due to the abolishment of the bacteriophage families *Myoviridae, Siphoviridae*, and *Podoviridae* in 2022, many bacteriophage sequences now find themselves without a corresponding viral family. This can be seen in deeper taxonomic assignments (Figure 2-B), with most sequences assigned to *Caudoviricetes* unable to be resolved any further – the sample with the highest RA of deeper *Caudoviricetes* taxonomy was August 2019 with only 18% assigned family-level taxonomy. Generally, those that were able to be assigned taxonomy past *Caudoviricetes* appeared to be ephemeral; viral families *Straboviridae* and *Kyanoviridae* were only found in three and four of the total samples respectively, while the order *Crassvirales* was found only in three of the samples. In addition, these families (and order, in the case of *Crassvirales*), were not particularly abundant, with the highest RA being *Crassvirales* in June 2019 (5%). *Ackermannviridae, Autographivirales, Demerecviridae*, and *Herelleviridae* were assigned at >1% RA and were included under “other taxa” (Figure 2-B). In comparison, most sequences of the class *Megaviricetes* were assigned a family-level taxonomy. The two main families were assigned as either *Phycodnaviridae* or *Mimiviridae*, viruses that infect primarily eukaryotic algae and protists, respectively. Both families were persistent across multiple samples, with higher RA in June 2019 and August 2019. Additionally, *Phycodnaviridae* and *Mimiviridae* showed a decline in RA in the winter sample of January 2020, indicating that viruses from *Megaviricetes* could have dynamic abundances that change with seasons.

One possible explanation for the differences seen between samples and across seasons may be the number of raw reads obtained after sequencing, resulting in a varied number of assembled contigs (Table S1). This could have potentially skewed our analysis and introduced a bias that would have omitted from detection certain families or their identified RA. However, rarefaction analysis (Figure S1) indicates that there was sufficient sequencing depth to cover the full extent of the cell-associated viral diversity in our samples. A more likely explanation is that the observed differences in RA and its diversity represent true temporal variations of the cell-associated virus community within BTL, which is indicative of its dynamic nature. Indeed, there are numerous factors that can affect the dynamic nature of a microbial community in a lake such as BTL. These include host range and the type of microbial host community, dissolved organic compound (DOC) composition, environmental conditions, dispersal mechanisms, seasonal patterns, evolutionary adaptations, infection dynamics (e.g., adsorption rates, burst sizes, and latent periods), resistance to environmental degradation, and competitive interactions between viruses over the same host for infection (Clokie *et al*., 2011; Mohiuddin and Schellhorn, 2015; Batinovic *et al*., 2019; Li *et al*., 2021; Sandor *et al*., 2026).

Finally, we also detected two virophage signatures (i.e., satellite viruses that co-infect with large algal viruses) belonging to the class *Virophaviricetes* (Figure 2-A), one in May 2019, and the other in June 2019. While virophages are still a relatively underrepresented group of viruses in databases, Fischer (2021) notes that the majority of virophages recovered from metagenomes so far are from lakes. It is therefore not surprising that we detect them in our metagenomic analysis. However, it is surprising that their RA was low in our study, given prior work that has shown that up to two thirds of the virus community in a lake may consist of virophages (Palermo *et al*., 2019). Virophages are mostly associated with *Mimiviridae* (Tokarz-Deptula *et al*., 2023), and while *Mimiviridae* are present within BTL, they do not appear to be particularly abundant in comparison to other viruses. We therefore conclude that protist viruses from *Mimiviridae* and virophages likely play a lesser role in BTL compared to in other aquatic habitats.

### 3.2 Host identification of the bacteriophage community

To gain insights about the hosts with which the Big Turkey Lake (BTL) cell-associated virus community is associated with, we ran all viral contigs (total of 998; Table S1) through the iPHoP pipeline, which is specifically designed to computationally predict bacterial and archaeal hosts from viral data (Roux, 2023). This analysis was able to predict hosts (Figure 2-C) for 21.8% of the viral contigs, the majority of which had an iPHoP confidence score of 90-96%.

Virus hosts belonging to the phyla *Actinobacteriota* and *Bacteriodota* were predicted in all samples across the different seasons. This is in accordance with prior findings that report on the presence of these phyla across those samples in BTL (Cameron *et al*., 2022; Cameron *et al*., 2024). Hosts belonging to the phylum Cyanobacteria were predicted for all samples apart from January 2020. This is in accordance with findings by Cameron *et al*. (2022) that show nearly complete absence of cyanobacteria in the ice-covered months of 2020, in both BTL and the rest of the lakes that make up the TLW.

Similarly, hosts belonging to the phylum *Pseudomonadota* were predicted in all samples apart from January 2020. A lack of *Pseudomonadota* host predictions for January is surprising, given that their RA in that sample was reported by Cameron *et al*. (2022) to be >50% of the total bacterial community. There are two possible explanations for this discrepancy. The first is that viruses are indeed associated with *Pseudomonadota* in the ice-covered winter months in BTL, but the iPHoP pipeline failed to predict hosts for the viral contigs in this sample given the low percentage of host predictions for identified viral contigs. The second is that *Pseudomonadota* in BTL is free of viral predation in the winter, and that their viruses are within the “free fraction” of the virus community (e.g., not cell associated/actively infecting).

The majority of host predictions belonging to the phylum *Verrucomicrobiota* were in August 2019 and January 2020, where they comprised 36% and 70% of the total predictions in those samples, respectively; however, *Verrucomicrobiota* were not assigned as viral hosts in either May 2019 or June 2019 despite this phylum being previously detected (Cameron et al., 2022). Although we cannot rule out active infection of *Verrucomicrobiota* entirely, this may suggest real temporal variation in BTL between actively infecting viruses associated with *Verrucomicrobiota* members in some of our samples, and those that are in the “free fraction” in the rest of our samples, which would not be detected based on our methods.

Finally, our host predictions were able to also provide new insights on the genus level. Some of the viral contigs in our July 2018 sample were predicted to infect members of the *Cyanobium* (unicellular cyanobacteria; iPHoP confidence score of 91-93), *Vulcanococcus* (cyanobacteria capable of forming microcolonies; iPHoP confidence score of 90-92.5), and *Tolypothrix* (filamentous cyanobacteria’; iPHoP confidence score of 90.3) genera. To the best of our knowledge this is the first report that identifies possible virus signatures associated with sequences attributed to these cyanobacterial genera.

### 3.3 Auxiliary metabolic genes

Viral hits to auxiliary metabolic genes (AMGs) were found within Big Turkey Lake (BTL) across all samples with seasonal variation in the number of AMGs found (Figure 3; Table S3). Of the total 88 genes identified as potential AMGs, 68 had at least one potential viral signature in the AMG flanking regions (Table S3). AMGs for nucleotide metabolism were the most abundant, with a combined total of 27 genes. Within those, there were multiple genes such as guanine reductase (*guaC*), guanylate kinase (*gmk*), and ribonucleoside-diphosphate reductase (*nrdA*) represented across multiple samples. AMGs in this category can bolster the biosynthesis of nucleic acids, which is an important step in viral replication (Gao *et al*., 2016; Crummett *et al*., 2016). In addition, there were also multiple AMGs involved in folate biosynthesis. This includes genes such as thymidylate synthase (*thyX, thyA*), and dihydrofolate reductase (*folA*), all of which have been found in viruses previously (Luo *et al*., 2022). Given that viruses downregulate production of host proteins (Fabricant & Kennel, 1970; Hurwitz & U’ren, 2016), and folate is an essential nutrient for DNA and protein synthesis (Bermingham & Derrick, 2002), viruses with those AMGs in their possession might be forcing their hosts to produce more folic acid to boost their own replication.

**Figure 3.**
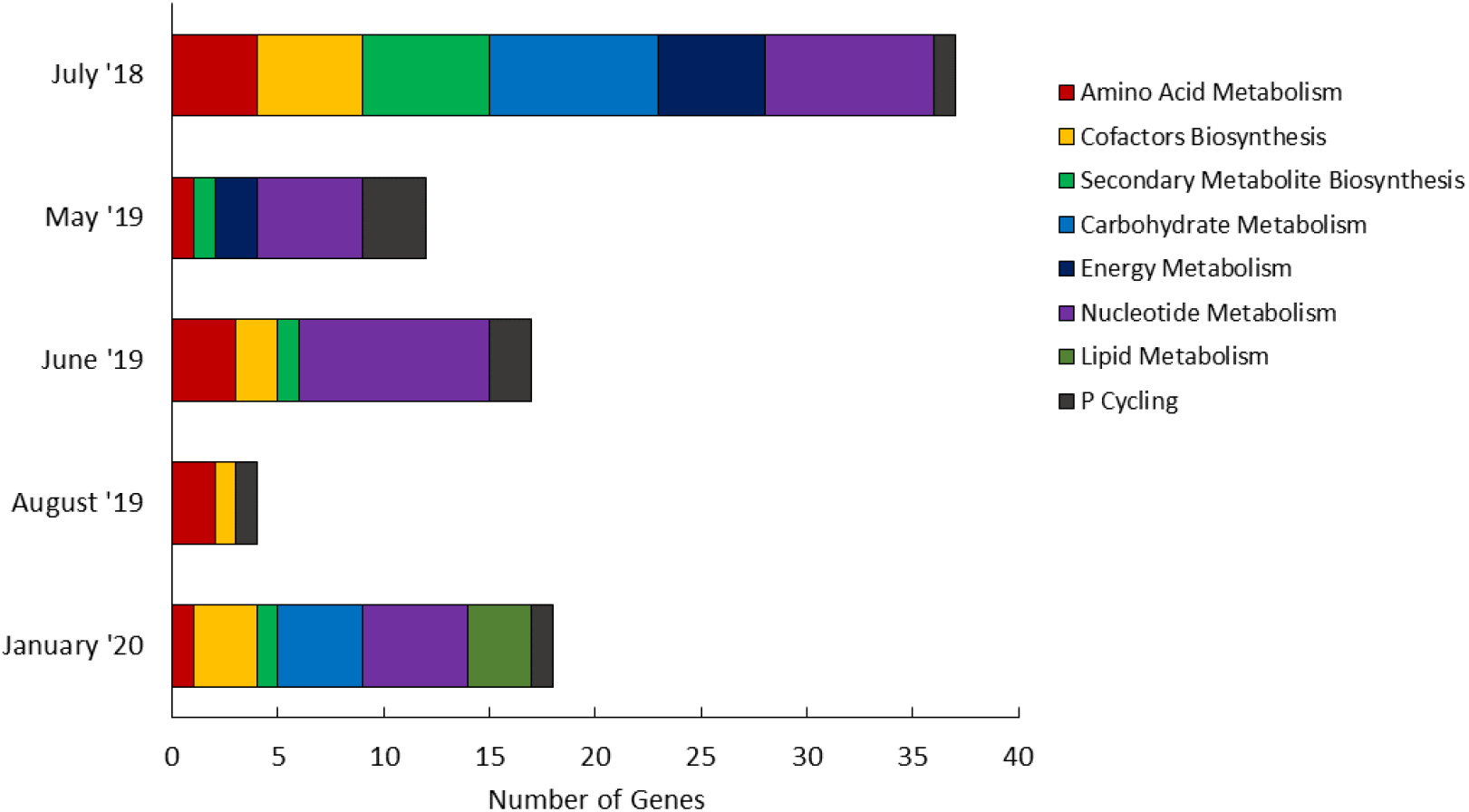
Auxiliary metabolic gene (AMG) profiles in Big Turkey Lake across samples/seasons and the number of identified genes within each AMG category.

Other key AMGs that we detected were those involved in energy metabolism pathways. Specifically, succinyl-CoA synthetase (*sucD*) and malate dehydrogenase (*mdh*) were represented across multiple samples, both of which are genes involved in the TCA cycle (Minàrik *et al*., 2002; Buck *et al*., 1985). Also, in the category of energy metabolism, we identified AMGs involved in photosynthesis: photosystem II (*psbK*). Oceanic phages were found to be common carriers of photosystem II genes, which makes them key players in global photosynthesis rates (Lindell *et al*., 2005; Heyerhoff *et al*., 2022). Metatranscriptomic studies show that viruses can contribute >40% of *psbA* expression within an environment (Sieradzki *et al*., 2019), making them key players in global oxygen production. Other AMGs identified included those involved in nitrogen fixation (*nifU*), found in June 2018 and May 2019. The collective presence of these AMGs in our study therefore indicates that they likely play an important role in primary productivity and nitrogen cycling in BTL.

Genes related to the phosphorus regulon (*phoH*) (Hsieh & Wanner, 2010) were found across all five samples, indicating viruses in BTL could play a role in phosphorus cycling. Given that BTL is classified as oligotrophic (Jeffries *et al*., 1988), and the presence of phosphorus-scavenging bacterial taxa have been previously identified (Cameron *et al*., 2024), the presence of genes related to phosphorus uptake have the potential to offer an evolutionary advantage for viruses that encode them. In addition, as *phoH* has been previously used as a marker for phage diversity in marine environments (Goldsmith *et al*., 2011), the ubiquity of *phoH* across all our samples suggests that this gene could be used as a similar diversity marker in BTL.

Further, consideration of the broader ecological context provides a basis for examining and integrating the viral community insights presented here within the TLW system. Changes in the composition of dissolved organic matter are increasingly identified as drivers of temporal dynamics in microbial community composition (Sandor *et al*., 2026). Previous work in TLW demonstrated that hydrological flow paths and event-driven connectivity strongly influence dissolved organic matter and phosphorus transport (Fines *et al*., 2023; Gray *et al*., 2024), with prior microbial community analyses repeatedly identifying the presence of phosphorus-scavenging taxa in this oligotrophic system (Cameron *et al*., 2024). Within this framework, the observed seasonal shifts in cell-associated viral communities and the ubiquitous detection of phosphorus-acquisition-related AMGs are consistent with the actively-infecting viral fraction being associated with phosphorus-efficient microbial hosts. Collectively, these findings suggest likely coupling between watershed-scale transport dynamics, seasonal microbial succession, and host-viral interactions within BTL, and potentially the TLW system as a whole.

Interestingly, AMGs involved in lipid metabolism were only detected in January 2020, making them the most infrequently found AMGs during the study period. These genes were 1,2-diacylglycerol beta-glucosyltransferase (*ugtP*) and 3-oxoacyl-[acyl-carrier protein] reductase and synthase (*fabF, fabG*), both of which are important in the biosynthesis of membrane lipids (Matsuoka *et al*., 2016; Guo *et al*., 2019). The *fabG* gene is highly conserved among bacterial species, emphasizing its importance (Guo et al., 2019). The detection of these AMGs in the winter is thus intriguing, given the fact that some bacteria adapt to cold temperatures by increasing production of membrane lipids and fatty acids (Hassan *et al*., 2020).

### 3.4 Viral metagenome-assembled genomes (vMAGs)

From our metagenomic data, we constructed 103 viral bins (representing a total of 360 viral contigs) that spanned across all five samples. Analysis of these bins by CheckV indicated that only six had a completeness of >90%, which made them candidates for vMAG consideration. These vMAG candidates had a genome size range of 11-45 kbp (Figure 4). Further analysis with the IMG/VR database revealed that all six vMAGs generated from BTL were aligned with *Caudoviricetes* phages (Table S4). In addition, four of the six vMAGs were able to be assigned host phyla using the iPHoP pipeline, with two being assigned to *Bacteriodota*, one to *Cyanobacteriota*, and one to *Pseudomonadota*.

**Figure 4.**
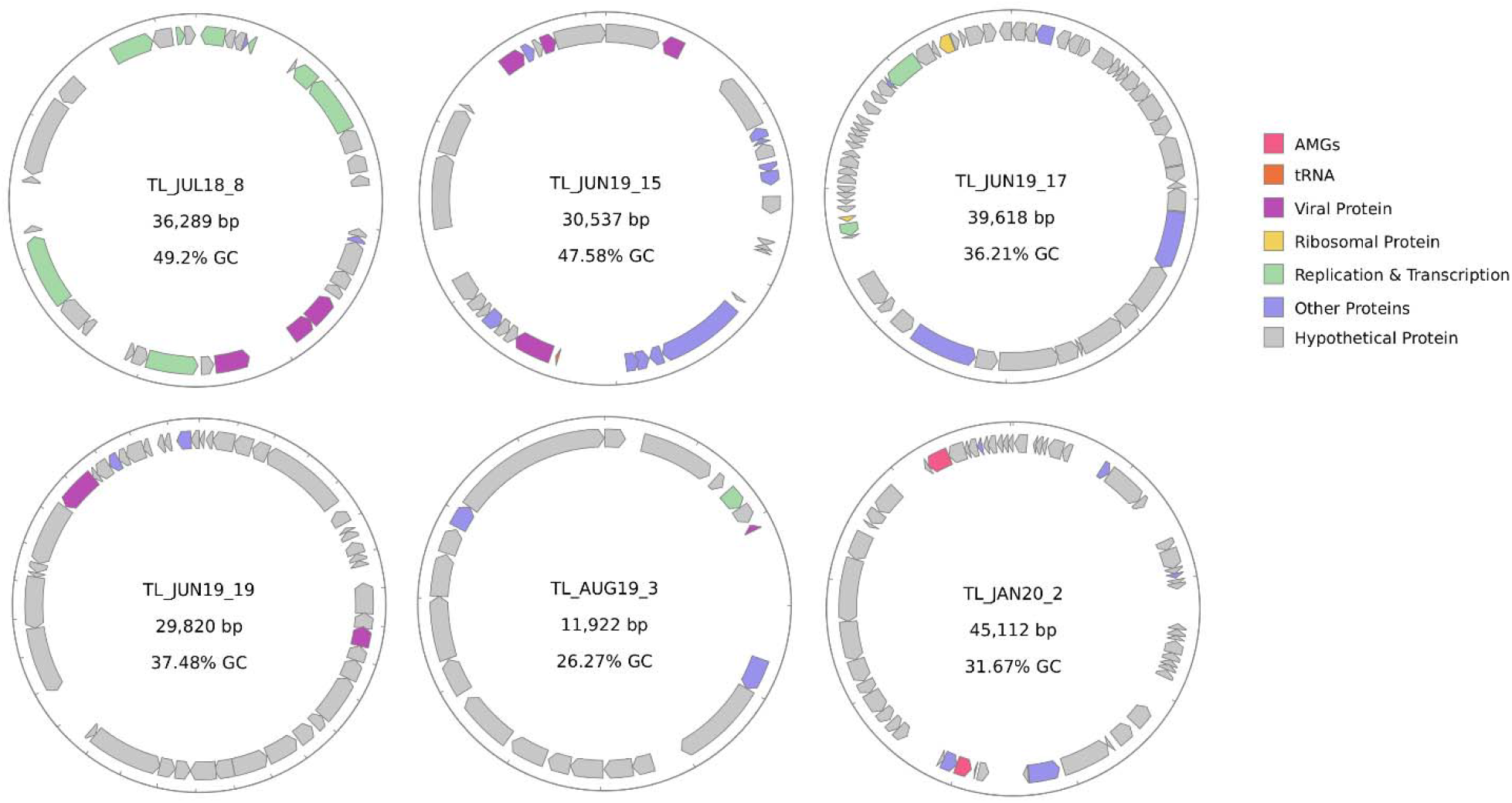
Viral Metagenome-Assembled Genomes (vMAGs) in Big Turkey Lake. vMAGs are circularized for visual clarity using *gbdraw* software. Colors represent structural viral proteins, other annotated proteins, hypothetical proteins (unknown function), tRNAs, virus proteins related to replication and transcription, AMGs, and ribosomal proteins (see Table S5 for more details). Note that although the latter can be considered too as AMGs, they are indicated here separately to highlight the significance of their detection.

From the annotated genes (Table S5), we found that two vMAGs (bin 8 and bin 15) contained coding regions for integrases, indicating a possible temperate lifecycle. Coding regions for tRNAs were also annotated in bin 2 and bin 15, for histidine and serine, respectively. Annotations in bin 17 included two proteins involved with ribosomes – specifically, ribosomal protein L18 and 23S rRNA/tRNA-pseudouridylate synthase. While the separation between viruses and other forms of life has been previously defined by the absence of ribosomes (Raoult & Forterre, 2008), bacteriophages have been found to encode ribosomal proteins, with the current highest number being six within one jumbophage (Meza-Padilla *et al*., 2026).

Currently, very few freshwater studies in Canada have reported on vMAGs (Mohiuddin & Schellhorn, 2015), leading to an under-representation of northern freshwater viruses. Assembly of novel viral genomes from uncultivated viruses is essential to increase our understanding of virus-host connections and their ecological implications (Nayfach *et al*., 2021).

## 4 Conclusions

Our work focused only on viruses associated with cells and did not include an analysis of “free living” viruses within the water column, or of those locked within the sediment of Big Turkey Lake. Nevertheless, it provides novel insights into the virus community that is likely actively infecting microbial hosts in this lake across seasons. We reveal i) a virus community dominated by *Caudoviricetes*, along with a large proportion of unclassified viruses, suggesting that many of the viruses in BTL are unique; ii) that the cell-associated virus community in BTL is dynamic between seasons and between years, and iii) that it is possible to assemble novel virus genomes even from small amounts of data, originating from a cellular microbial fraction.

We also illustrate the power of machine learning tools such as iPHoP for predicting the hosts that classified and unclassified viruses are expected to be actively infecting in a setting such as BTL. In our work, this tool enabled a glimpse into the identity of the microbial hosts that were likely being infected in different seasons, a critical aspect in improving our understanding of the role of viruses in freshwater. Importantly, this tool allowed us to report for the first time on potential cyanophage associations to *Cyanobium, Vulcanococcus*, and *Tolypothrix*.

Finally, we report on functionally and ecologically important viral genes such as AMGs and hypothesize on their potential role within BTL. Based on the limited data that we have, it is difficult to implicate which specific ecological, biological and physicochemical parameters might be driving their distribution or influence the AMG profile across our sampled seasons. Such profiles are driven predominantly by viral lifestyle (lytic vs lysogenic), habitat characteristics, and host identities (Luo *et al*., 2022); all aspects that should be included in future investigations.

## 5 Conflict of Interest

The authors declare that the research was conducted in the absence of any commercial or financial relationships that could be construed as a potential conflict of interest.

## 6 Author Contributions

The work was conceptualized by C.C.C. and J.I.N.; Samples were collected by E.S.C.; Access to samples was allowed by K.M.M. and M.B.E. through the forWater network (https://www.forwater.ca/); data analysis and interpretation was performed by C.C.C., N.T. and J.I.N.; original draft preparation of the manuscript was done by C.C.C and J.I.N.; review and editing was performed by C.C.C, N.T., E.S.C, M.B.E. and J.I.N.; student supervision and project administration were done by J.I.N.; and funding was acquired by J.I.N. All authors have read and agreed to the published version of the manuscript.

## 7 Funding

This work was supported by Natural Sciences and Engineering Research Council of Canada (RGPIN-2022-03350 and DGECR-2022-00329) and a 2021 Phycological Society of America Norma J. Lang Early Career Research Fellowship, awarded to J.I.N.

## 8 Acknowledgments

We acknowledge the support of the forWater NSERC Network for Forested Drinking Water Source Protection Technologies [NETGP-494312-16] for allowing us access to sampling site and samples, and the Digital Research Alliance of Canada, which allowed for additional supported computing, storage, or data services. We would like to also thank Mr. Isaac Meza-Padilla for helping us create the map in Figure 1.

## 10 Data Availability Statement

The metagenomic reads from this study can be found in the SRA under the accession number PRJNA1018242. Accession numbers for the generated vMAGs are PP191267, PP191268, PP191269, PP191270, PP191271, and PP191272.

## Notes

### Competing Interest Statement

The authors have declared no competing interest.

### Summary of Updates

Author name edits - manuscript unchanged.

