## Supplemental Tables for "Cell-associated viral community composition and its functional potential in a dimictic lake on the Canadian Shield"

### Supplementary

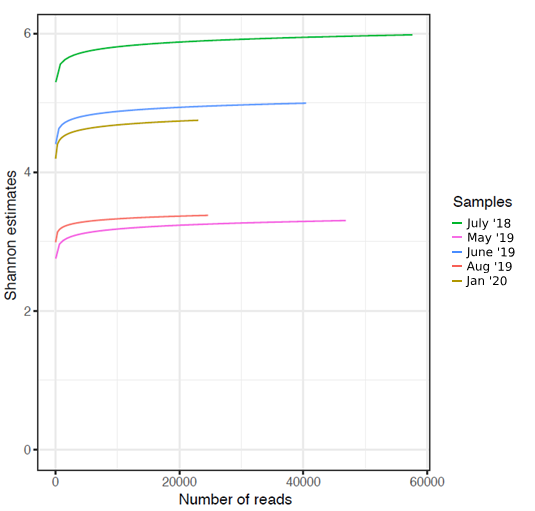

**Figure S1.** Rarefaction curves on viral community of Big Turkey Lake samples, showing Shannon diversity index at various sequencing depths.

**Table S1.** Sample Summary from Big Turkey Lake. Samples from 2018 were taken from 1 meter below Secchi depth, while those sampled from 2019 were taken at Secchi depth. *Due to ice cover at the time of sampling, the January 2020 sample was taken at surface level.

| **Sample** | **Date** | **Depth** | **Air**  **°C** | **DNA concentration** | **# of reads (M)** | **total assembled contigs** | **viral contigs** | **% viral contigs** |
| --- | --- | --- | --- | --- | --- | --- | --- | --- |
| July 2018 | 18/07/2018 | 8 | 23.4 | 11.7 | 4.54 | 82018 | 581 | 0.708 |
| May 2019 | 23/05/2019 | 5.25 | N/A | 1.6 | 3.75 | 15221 | 110 | 0.723 |
| June 2019 | 28/06/2019 | 5.75 | 19.3 | 5.1 | 2.8 | 23257 | 135 | 0.58 |
| August 2019 | 21/07/2019 | 5 | 21.2 | 4.9 | 3.52 | 19214 | 42 | 0.219 |
| January 2020 | 23/01/2020 | 0* | N/A | 3.5 | 3.8 | 20765 | 130 | 0.626 |

**Table S2.** Quality control of sequncind samples from Big Turkey Lake was performed using the Metagenome-Atlas pipeline.

| **Sample** | **Reads** | | **Bases** | |
| --- | --- | --- | --- | --- |
|  | **Raw** | **After QC** | **Raw** | **After QC** |
| July 2018 | 4.54M | 4.51M | 2.73B | 2.34B |
| May 2019 | 3.75M | 3.61M | 1.45B | 1.41B |
| June 2019 | 2.80M | 2.71M | 1.12B | 1.09B |
| August 2019 | 3.51M | 3.27M | 1.14B | 1.09B |
| January 2020 | 3.8M | 3.67M | 1.36B | 1.34B |

**Table S3.** Auxiliary metabolic genes (AMGs) in Big Turkey Lake. Pathways of designated AMGs were assigned based on the KEGG Orthology database. Viral flanking regions were defined as regions surrounding the gene in question with either known viral genes, or a region of very short, hypothetical proteins.

| **Sample** | **Name** | **Definition** | **Pathway** | **# of potential viral flanking regions** |
| --- | --- | --- | --- | --- |
| TL_JUL18 | speD, AMD1 | S-adenosylmethionine decarboxylase [EC:4.1.1.50] | Amino acid metabolism | 1 |
| TL_JUL18 | glnA, GLUL | glutamine synthetase [EC:6.3.1.2] | Amino acid metabolism | 1 |
| TL_JUL18 | DNMT1, dcm | DNA (cytosine-5)-methyltransferase 1 [EC:2.1.1.37] | Amino Acid Metabolism | 2 |
| TL_JUL18 | mhpC | 2-hydroxy-6-oxonona-2,4-dienedioate hydrolase [EC:3.7.1.14] | Amino acid metabolism | 1 |
| TL_JUL18 | DHFR, folA | dihydrofolate reductase [EC:1.5.1.3] | Biosynthesis of cofactors | 2 |
| TL_JUL18 | pyrF | orotidine-5'-phosphate decarboxylase [EC:4.1.1.23] | Biosynthesis of cofactors | 2 |
| TL_JUL18 | hemH, FECH | protoporphyrin/coproporphyrin ferrochelatase [EC:4.99.1.1 4.99.1.9] | Biosynthesis of cofactors | 0 |
| TL_JUL18 | iscS, NFS1 | cysteine desulfurase [EC:2.8.1.7] | Biosynthesis of cofactors | 0 |
| TL_JUL18 | GAE, cap1J | UDP-glucuronate 4-epimerase [EC:5.1.3.6] | Biosynthesis of cofactors | 0 |
| TL_JUL18 | rfbD, rmlD | dTDP-4-dehydrorhamnose reductase [EC:1.1.1.133] | Biosynthesis of secondary metabolites | 2 |
| TL_JUL18 | rfbA, rmlA, rffH | glucose-1-phosphate thymidylyltransferase [EC:2.7.7.24] | Biosynthesis of secondary metabolites | 2 |
| TL_JUL18 | trpC | indole-3-glycerol phosphate synthase [EC:4.1.1.48] | Biosynthesis of secondary metabolites | 0 |
| TL_JUL18 | E2.2.1.6L, ilvB, ilvG, ilvI | acetolactate synthase I/II/III large subunit [EC:2.2.1.6] | Biosynthesis of secondary metabolites | 0 |
| TL_JUL18 | rfbB, rmlB, rffG | dTDP-glucose 4,6-dehydratase [EC:4.2.1.46] | Biosynthesis of secondary metabolites | 2 |
| TL_JUL18 | rfbC, rmlC | dTDP-4-dehydrorhamnose 3,5-epimerase [EC:5.1.3.13] | Biosynthesis of secondary metabolites | 2 |
| TL_JUL18 | gmd, GMDS | GDPmannose 4,6-dehydratase [EC:4.2.1.47] | Carbohydrate Metabolism | 1 |
| TL_JUL18 | gmd, GMDS | GDPmannose 4,6-dehydratase [EC:4.2.1.47] | Carbohydrate Metabolism | 0 |
| TL_JUL18 | gmd, GMDS | GDPmannose 4,6-dehydratase [EC:4.2.1.47] | Carbohydrate Metabolism | 1 |
| TL_JUL18 | galE, GALE | UDP-glucose 4-epimerase [EC:5.1.3.2] | Carbohydrate Metabolism | 1 |
| TL_JUL18 | pgmB | beta-phosphoglucomutase [EC:5.4.2.6] | Carbohydrate Metabolism | 0 |
| TL_JUL18 | TSTA3, fcl | GDP-L-fucose synthase [EC:1.1.1.271] | Carbohydrate Metabolism | 1 |
| TL_JUL18 | TSTA3, fcl | GDP-L-fucose synthase [EC:1.1.1.271] | Carbohydrate Metabolism | 0 |
| TL_JUL18 | glmS, GFPT | glutamine---fructose-6-phosphate transaminase (isomerizing) [EC:2.6.1.16] | Carbohydrate Metabolism | 1 |
| TL_JUL18 | iscU, nifU | nitrogen fixation protein NifU and related proteins | Energy Metabolism | 0 |
| TL_JUL18 | mdh | malate dehydrogenase [EC:1.1.1.37] | Energy Metabolism | 0 |
| TL_JUL18 | sucD | succinyl-CoA synthetase alpha subunit [EC:6.2.1.5] | Energy Metabolism | 1 |
| TL_JUL18 | psbK | photosystem II PsbK protein | Energy Metabolism | 1 |
| TL_JUL18 | ndhB | NAD(P)H-quinone oxidoreductase subunit 2 [EC:7.1.1.2] | Energy Metabolism | 0 |
| TL_JUL18 | E1.7.1.7, guaC | GMP reductase [EC:1.7.1.7] | Nucleotide metabolism | 1 |
| TL_JUL18 | thyA, TYMS | thymidylate synthase [EC:2.1.1.45] | Nucleotide metabolism | 1 |
| TL_JUL18 | APRT, apt | adenine phosphoribosyltransferase [EC:2.4.2.7] | Nucleotide metabolism | 0 |
| TL_JUL18 | E2.7.4.8, gmk | guanylate kinase [EC:2.7.4.8] | Nucleotide metabolism | 0 |
| TL_JUL18 | dcd | dCTP deaminase [EC:3.5.4.13] | Nucleotide metabolism | 2 |
| TL_JUL18 | dut, DUT | dUTP pyrophosphatase [EC:3.6.1.23] | Nucleotide metabolism | 1 |
| TL_JUL18 | pyrG, CTPS | CTP synthase [EC:6.3.4.2] | Nucleotide metabolism | 2 |
| TL_JUL18 | thyX, thy1 | thymidylate synthase (FAD) [EC:2.1.1.148] | Nucleotide metabolism | 1 |
| TL_JUL18 | PhoH | phosphate starvation-inducible protein PhoH | Phosphorus Cycling | 1 |
| TL_MAY19 | rtpR | ribonucleoside-triphosphate reductase (thioredoxin) [EC:1.17.4.2] | Nucleotide Metabolism | 2 |
| TL_MAY19 | dut, DUT | dUTP pyrophosphatase [EC:3.6.1.23] | Nucleotide Metabolism | 1 |
| TL_MAY19 | E4.6.1.1 | adenylate cyclase [EC:4.6.1.1] | Nucleotide Metabolism | 1 |
| TL_MAY19 | thyX, thy1 | thymidylate synthase (FAD) [EC:2.1.1.148] | Nucleotide Metabolism | 1 |
| TL_MAY19 | HDDC3 | guanosine-3',5'-bis(diphosphate) 3'-pyrophosphohydrolase [EC:3.1.7.2] | Nucleotide Metabolism | 1 |
| TL_MAY19 | iscU, nifU | nitrogen fixation protein NifU and related proteins | Energy Metabolism | 2 |
| TL_MAY19 | erpA | iron-sulfur cluster insertion protein | Energy Metabolism | 2 |
| TL_MAY19 | K23144 | glucosamine-1-phosphate N-acetyltransferase | Biosynthesis of Secondary Metabolites | 1 |
| TL_MAY19 | map | methionyl aminopeptidase [EC:3.4.11.18] | Amino Acid Metabolism | 0 |
| TL_MAY19 | PhoH | phosphate starvation-inducible protein PhoH | Phosphorus Cycling | 1 |
| TL_MAY19 | PhoH | phosphate starvation-inducible protein PhoH | Phosphorus Cycling | 1 |
| TL_MAY19 | PhoH | phosphate starvation-inducible protein PhoH | Phosphorus Cycling | 1 |
| TL_JUN19 | E1.7.1.7, guaC | GMP reductase [EC:1.7.1.7] | Nucleotide Metabolism | 1 |
| TL_JUN19 | E1.17.4.1A, nrdA, nrdE | ribonucleoside-diphosphate reductase alpha chain [EC:1.17.4.1] | Nucleotide Metabolism | 1 |
| TL_JUN19 | E1.17.4.1A, nrdA, nrdE | ribonucleoside-diphosphate reductase alpha chain [EC:1.17.4.1] | Nucleotide Metabolism | 2 |
| TL_JUN19 | rtpR | ribonucleoside-triphosphate reductase (thioredoxin) [EC:1.17.4.2] | Nucleotide Metabolism | 1 |
| TL_JUN19 | thyA, TYMS | thymidylate synthase [EC:2.1.1.45] | Nucleotide Metabolism | 1 |
| TL_JUN19 | tdk, TK | thymidine kinase [EC:2.7.1.21] | Nucleotide Metabolism | 1 |
| TL_JUN19 | tdk, TK | thymidine kinase [EC:2.7.1.21] | Nucleotide Metabolism | 1 |
| TL_JUN19 | thyX, thy1 | thymidylate synthase (FAD) [EC:2.1.1.148] | Nucleotide Metabolism | 1 |
| TL_JUN19 | thyX, thy1 | thymidylate synthase (FAD) [EC:2.1.1.148] | Nucleotide Metabolism | 1 |
| TL_JUN19 | acpP | acyl carrier protein | Biosynthesis of Secondary Metabolites | 1 |
| TL_JUN19 | pdxA | 4-hydroxythreonine-4-phosphate dehydrogenase [EC:1.1.1.262] | Biosynthesis of cofactors | 1 |
| TL_JUN19 | gpt | xanthine phosphoribosyltransferase [EC:2.4.2.22] | Biosynthesis of cofactors | 1 |
| TL_JUN19 | trpC | indole-3-glycerol phosphate synthase [EC:4.1.1.48] | Amino Acid Metabolism | 1 |
| TL_JUN19 | speD, AMD1 | S-adenosylmethionine decarboxylase [EC:4.1.1.50] | Amino Acid Metabolism | 2 |
| TL_JUN19 | speD, AMD1 | S-adenosylmethionine decarboxylase [EC:4.1.1.50] | Amino Acid Metabolism | 0 |
| TL_JUN19 | PhoH | phosphate starvation-inducible protein PhoH | Phosphorus Cycling | 1 |
| TL_JUN19 | PhoH | phosphate starvation-inducible protein PhoH | Phosphorus Cycling | 2 |
| TL_AUG19 | DNMT1, dcm | DNA (cytosine-5)-methyltransferase 1 [EC:2.1.1.37] | Amino Acid Metabolism | 1 |
| TL_AUG19 | glmS, GFPT | glutamine---fructose-6-phosphate transaminase (isomerizing) [EC:2.6.1.16] | Amino Acid Metabolism | 1 |
| TL_AUG19 | queF | 7-cyano-7-deazaguanine reductase [EC:1.7.1.13] | Biosynthesis of cofactors | 1 |
| TL_AUG19 | PhoH | phosphate starvation-inducible protein PhoH | Phosphorus Cycling | 1 |
| TL_JAN20 | E1.17.4.1A, nrdA, nrdE | ribonucleoside-diphosphate reductase alpha chain [EC:1.17.4.1] | Nucleotide Metabolism | 1 |
| TL_JAN20 | E1.17.4.1A, nrdA, nrdE | ribonucleoside-diphosphate reductase alpha chain [EC:1.17.4.1] | Nucleotide Metabolism | 1 |
| TL_JAN20 | E1.17.4.1B, nrdB, nrdF | ribonucleoside-diphosphate reductase beta chain [EC:1.17.4.1] | Nucleotide Metabolism | 1 |
| TL_JAN20 | comEB | dCMP deaminase [EC:3.5.4.12] | Nucleotide Metabolism | 2 |
| TL_JAN20 | dut, DUT | dUTP pyrophosphatase [EC:3.6.1.23] | Nucleotide Metabolism | 2 |
| TL_JAN20 | fabG, OAR1 | 3-oxoacyl-[acyl-carrier protein] reductase [EC:1.1.1.100] | Lipid Metabolism | 0 |
| TL_JAN20 | ugtP | processive 1,2-diacylglycerol beta-glucosyltransferase [EC:2.4.1.315] | Lipid Metabolism | 2 |
| TL_JAN20 | fabF, OXSM, CEM1 | 3-oxoacyl-[acyl-carrier-protein] synthase II [EC:2.3.1.179] | Lipid Metabolism | 0 |
| TL_JAN20 | glmS, GFPT | glutamine---fructose-6-phosphate transaminase (isomerizing) [EC:2.6.1.16] | Carbohydrate Metabolism | 1 |
| TL_JAN20 | gmd, GMDS | GDPmannose 4,6-dehydratase [EC:4.2.1.47] | Carbohydrate Metabolism | 0 |
| TL_JAN20 | TSTA3, fcl | GDP-L-fucose synthase [EC:1.1.1.271] | Carbohydrate Metabolism | 2 |
| TL_JAN20 | UXS1, uxs | UDP-glucuronate decarboxylase [EC:4.1.1.35] | Carbohydrate Metabolism | 0 |
| TL_JAN20 | acpP | acyl carrier protein | Biosynthesis of Secondary Metabolites | 0 |
| TL_JAN20 | GCH1, folE | GTP cyclohydrolase IA [EC:3.5.4.16] | Biosynthesis of Cofactors | 1 |
| TL_JAN20 | queD, ptpS, PTS | 6-pyruvoyltetrahydropterin/6-carboxytetrahydropterin synthase [EC:4.2.3.12 4.1.2.50] | Biosynthesis of Cofactors | 1 |
| TL_JAN20 | ribBA | 3,4-dihydroxy 2-butanone 4-phosphate synthase / GTP cyclohydrolase II [EC:4.1.99.12 3.5.4.25] | Biosynthesis of Cofactors | 1 |
| TL_JAN20 | dapB | 4-hydroxy-tetrahydrodipicolinate reductase [EC:1.17.1.8] | Amino Acid Metabolism | 1 |
| TL_JAN20 | PhoH | phosphate starvation-inducible protein PhoH | Phosphorus Cycling | 1 |

**Table S4**. Big Turkey Lake viral metagenome assembled Genome (vMAG) summary and host prediction. Only vMAGs with a predicted completeness of >90% by CheckV were designated a ‘High-Quality’ vMAGs and used for further analysis.

| **vMAG** | **Sample** | **Predicted Completeness** | **Viral Class** | **iPHoP predicted host phyla** |
| --- | --- | --- | --- | --- |
| 8 | July 2018 | 100 | Caudoviricetes | Pseudomonadota |
| 15 | June 2019 | 92.64 | Caudoviricetes | Cyanobacteriota |
| 17 | June 2019 | 100 | Caudoviricetes | unassigned |
| 19 | June 2019 | 100 | Caudoviricetes | unassigned |
| 3 | August 2019 | 93.94 | Caudoviricetes | Bacteroidota |
| 2 | January 2020 | 100 | Caudoviricetes | Bacteroidota |

**Table S5.** Predicted protein annotations of vMAGs from Big Turkey Lake. Annotations from each vMAG constructed from the samples, as annotated by IMG/ER.

| **Sample and bin** | **protein coding genes** |
| --- | --- |
| TL_JUL18_8 | DNA-directed RNA polymerase subunit RPC12/RpoP |
| TL_JUL18_8 | hypothetical protein |
| TL_JUL18_8 | hypothetical protein |
| TL_JUL18_8 | nitrate/nitrite-specific signal transduction histidine kinase |
| TL_JUL18_8 | transcriptional regulator with XRE-family HTH domain |
| TL_JUL18_8 | hypothetical protein |
| TL_JUL18_8 | exodeoxyribonuclease-3 |
| TL_JUL18_8 | ATP-dependent DNA ligase |
| TL_JUL18_8 | hypothetical protein |
| TL_JUL18_8 | hypothetical protein |
| TL_JUL18_8 | hypothetical protein |
| TL_JUL18_8 | hypothetical protein |
| TL_JUL18_8 | uncharacterized membrane protein YeaQ/YmgE |
| TL_JUL18_8 | hypothetical protein |
| TL_JUL18_8 | hypothetical protein |
| TL_JUL18_8 | hypothetical protein |
| TL_JUL18_8 | HK97 family phage prohead protease |
| TL_JUL18_8 | HK97 family phage portal protein |
| TL_JUL18_8 | integrase |
| TL_JUL18_8 | hypothetical protein |
| TL_JUL18_8 | DNA polymerase-1 |
| TL_JUL18_8 | hypothetical protein |
| TL_JUL18_8 | hypothetical protein |
| TL_JUL18_8 | hypothetical protein |
| TL_JUL18_8 | hypothetical protein |
| TL_JUL18_8 | superfamily II DNA or RNA helicase |
| TL_JUL18_8 | hypothetical protein |
| TL_JUL18_8 | hypothetical protein |
| TL_JUL18_8 | hypothetical protein |
| TL_JUL18_8 | hypothetical protein |
| TL_JUL18_8 | superfamily II DNA or RNA helicase |
| TL_JUL18_8 | hypothetical protein |
| TL_JUL18_8 | predicted transcriptional regulator |
| TL_JUL18_8 | hypothetical protein |
| TL_JUN19_15 | hypothetical protein |
| TL_JUN19_15 | muramidase (phage lysozyme) |
| TL_JUN19_15 | hypothetical protein |
| TL_JUN19_15 | probable addiction module antidote protein |
| TL_JUN19_15 | putative component of toxin-antitoxin plasmid stabilization module |
| TL_JUN19_15 | hypothetical protein |
| TL_JUN19_15 | AbrB family looped-hinge helix DNA binding protein |
| TL_JUN19_15 | predicted nucleic acid-binding protein |
| TL_JUN19_15 | hypothetical protein |
| TL_JUN19_15 | hypothetical protein |
| TL_JUN19_15 | hypothetical protein |
| TL_JUN19_15 | hypothetical protein |
| TL_JUN19_15 | hypothetical protein |
| TL_JUN19_15 | type I restriction enzyme R subunit |
| TL_JUN19_15 | clan AA aspartic protease |
| TL_JUN19_15 | predicted nucleotidyltransferase |
| TL_JUN19_15 | HEPN domain-containing protein |
| TL_JUN19_15 | tRNA_Ser_TGA |
| TL_JUN19_15 | integrase |
| TL_JUN19_15 | hypothetical protein |
| TL_JUN19_15 | hypothetical protein |
| TL_JUN19_15 | single-strand DNA-binding protein |
| TL_JUN19_15 | hypothetical protein |
| TL_JUN19_15 | hypothetical protein |
| TL_JUN19_15 | hypothetical protein |
| TL_JUN19_15 | hypothetical protein |
| TL_JUN19_15 | hypothetical protein |
| TL_JUN19_15 | hypothetical protein |
| TL_JUN19_15 | uncharacterized protein involved in type VI secretion and phage assembly |
| TL_JUN19_15 | uncharacterized Zn-binding protein involved in type VI secretion |
| TL_JUN19_15 | hypothetical protein |
| TL_JUN19_15 | phage baseplate assembly protein W |
| TL_JUN19_15 | hypothetical protein |
| TL_JUN19_17 | hypothetical protein |
| TL_JUN19_17 | hypothetical protein |
| TL_JUN19_17 | GR25 family glycosyltransferase involved in LPS biosynthesis |
| TL_JUN19_17 | hypothetical protein |
| TL_JUN19_17 | hypothetical protein |
| TL_JUN19_17 | hypothetical protein |
| TL_JUN19_17 | hypothetical protein |
| TL_JUN19_17 | hypothetical protein |
| TL_JUN19_17 | hypothetical protein |
| TL_JUN19_17 | hypothetical protein |
| TL_JUN19_17 | hypothetical protein |
| TL_JUN19_17 | hypothetical protein |
| TL_JUN19_17 | hypothetical protein |
| TL_JUN19_17 | hypothetical protein |
| TL_JUN19_17 | hypothetical protein |
| TL_JUN19_17 | hypothetical protein |
| TL_JUN19_17 | hypothetical protein |
| TL_JUN19_17 | hypothetical protein |
| TL_JUN19_17 | DNA modification methylase |
| TL_JUN19_17 | hypothetical protein |
| TL_JUN19_17 | hypothetical protein |
| TL_JUN19_17 | hypothetical protein |
| TL_JUN19_17 | hypothetical protein |
| TL_JUN19_17 | hypothetical protein |
| TL_JUN19_17 | hypothetical protein |
| TL_JUN19_17 | hypothetical protein |
| TL_JUN19_17 | tape measure domain-containing protein |
| TL_JUN19_17 | hypothetical protein |
| TL_JUN19_17 | hypothetical protein |
| TL_JUN19_17 | hypothetical protein |
| TL_JUN19_17 | hypothetical protein |
| TL_JUN19_17 | transcriptional regulator with XRE-family HTH domain |
| TL_JUN19_17 | ribosomal protein L18 |
| TL_JUN19_17 | hypothetical protein |
| TL_JUN19_17 | hypothetical protein |
| TL_JUN19_17 | hypothetical protein |
| TL_JUN19_17 | hypothetical protein |
| TL_JUN19_17 | hypothetical protein |
| TL_JUN19_17 | hypothetical protein |
| TL_JUN19_17 | hypothetical protein |
| TL_JUN19_17 | hypothetical protein |
| TL_JUN19_17 | hypothetical protein |
| TL_JUN19_17 | hypothetical protein |
| TL_JUN19_17 | hypothetical protein |
| TL_JUN19_17 | hypothetical protein |
| TL_JUN19_17 | hypothetical protein |
| TL_JUN19_17 | hypothetical protein |
| TL_JUN19_17 | hypothetical protein |
| TL_JUN19_17 | hypothetical protein |
| TL_JUN19_17 | transcription elongation factor GreA-like protein |
| TL_JUN19_17 | replicative DNA helicase |
| TL_JUN19_17 | hypothetical protein |
| TL_JUN19_17 | hypothetical protein |
| TL_JUN19_17 | 23S rRNA-/tRNA-specific pseudouridylate synthase |
| TL_JUN19_17 | hypothetical protein |
| TL_JUN19_17 | hypothetical protein |
| TL_JUN19_17 | hypothetical protein |
| TL_JUN19_17 | hypothetical protein |
| TL_JUN19_17 | hypothetical protein |
| TL_JUN19_19 | hypothetical protein |
| TL_JUN19_19 | hypothetical protein |
| TL_JUN19_19 | hypothetical protein |
| TL_JUN19_19 | hypothetical protein |
| TL_JUN19_19 | hypothetical protein |
| TL_JUN19_19 | hypothetical protein |
| TL_JUN19_19 | hypothetical protein |
| TL_JUN19_19 | hypothetical protein |
| TL_JUN19_19 | hypothetical protein |
| TL_JUN19_19 | hypothetical protein |
| TL_JUN19_19 | hypothetical protein |
| TL_JUN19_19 | hypothetical protein |
| TL_JUN19_19 | hypothetical protein |
| TL_JUN19_19 | hypothetical protein |
| TL_JUN19_19 | lysozyme family protein |
| TL_JUN19_19 | hypothetical protein |
| TL_JUN19_19 | hypothetical protein |
| TL_JUN19_19 | hypothetical protein |
| TL_JUN19_19 | hypothetical protein |
| TL_JUN19_19 | hypothetical protein |
| TL_JUN19_19 | hypothetical protein |
| TL_JUN19_19 | hypothetical protein |
| TL_JUN19_19 | hypothetical protein |
| TL_JUN19_19 | hypothetical protein |
| TL_JUN19_19 | hypothetical protein |
| TL_JUN19_19 | hypothetical protein |
| TL_JUN19_19 | hypothetical protein |
| TL_JUN19_19 | hypothetical protein |
| TL_JUN19_19 | hypothetical protein |
| TL_JUN19_19 | hypothetical protein |
| TL_JUN19_19 | hypothetical protein |
| TL_JUN19_19 | hypothetical protein |
| TL_JUN19_19 | hypothetical protein |
| TL_JUN19_19 | phage terminase large subunit |
| TL_JUN19_19 | hypothetical protein |
| TL_JUN19_19 | hypothetical protein |
| TL_JUN19_19 | hydroxymethylpyrimidine pyrophosphatase-like HAD family hydrolase |
| TL_JUN19_19 | hypothetical protein |
| TL_JUN19_19 | hypothetical protein |
| TL_JUN19_19 | hypothetical protein |
| TL_JUN19_19 | hypothetical protein |
| TL_JUN19_19 | hypothetical protein |
| TL_JUN19_19 | single-strand DNA-binding protein |
| TL_JUN19_19 | hypothetical protein |
| TL_AUG19_3 | hypothetical protein |
| TL_AUG19_3 | hypothetical protein |
| TL_AUG19_3 | hypothetical protein |
| TL_AUG19_3 | predicted HTH transcriptional regulator |
| TL_AUG19_3 | hypothetical protein |
| TL_AUG19_3 | GH24 family phage-related lysozyme (muramidase) |
| TL_AUG19_3 | LEA14-like dessication related protein |
| TL_AUG19_3 | hypothetical protein |
| TL_AUG19_3 | hypothetical protein |
| TL_AUG19_3 | hypothetical protein |
| TL_AUG19_3 | hypothetical protein |
| TL_AUG19_3 | hypothetical protein |
| TL_AUG19_3 | hypothetical protein |
| TL_AUG19_3 | hypothetical protein |
| TL_AUG19_3 | hypothetical protein |
| TL_AUG19_3 | hypothetical protein |
| TL_AUG19_3 | hypothetical protein |
| TL_AUG19_3 | hypothetical protein |
| TL_AUG19_3 | ATP-dependent Lon protease |
| TL_AUG19_3 | hypothetical protein |
| TL_JAN20_2 | hypothetical protein |
| TL_JAN20_2 | hypothetical protein |
| TL_JAN20_2 | hypothetical protein |
| TL_JAN20_2 | hypothetical protein |
| TL_JAN20_2 | hypothetical protein |
| TL_JAN20_2 | hypothetical protein |
| TL_JAN20_2 | N-acetyl-anhydromuramyl-L-alanine amidase AmpD |
| TL_JAN20_2 | hypothetical protein |
| TL_JAN20_2 | hypothetical protein |
| TL_JAN20_2 | hypothetical protein |
| TL_JAN20_2 | hypothetical protein |
| TL_JAN20_2 | hypothetical protein |
| TL_JAN20_2 | uncharacterized membrane protein |
| TL_JAN20_2 | hypothetical protein |
| TL_JAN20_2 | hypothetical protein |
| TL_JAN20_2 | hypothetical protein |
| TL_JAN20_2 | hypothetical protein |
| TL_JAN20_2 | hypothetical protein |
| TL_JAN20_2 | hypothetical protein |
| TL_JAN20_2 | hypothetical protein |
| TL_JAN20_2 | hypothetical protein |
| TL_JAN20_2 | hypothetical protein |
| TL_JAN20_2 | hypothetical protein |
| TL_JAN20_2 | hypothetical protein |
| TL_JAN20_2 | hypothetical protein |
| TL_JAN20_2 | hypothetical protein |
| TL_JAN20_2 | hypothetical protein |
| TL_JAN20_2 | CubicO group peptidase |
| TL_JAN20_2 | hypothetical protein |
| TL_JAN20_2 | hypothetical protein |
| TL_JAN20_2 | tRNA_His_GTG |
| TL_JAN20_2 | GTP cyclohydrolase I |
| TL_JAN20_2 | 6-pyruvoyltetrahydropterin/6-carboxytetrahydropterin synthase |
| TL_JAN20_2 | hypothetical protein |
| TL_JAN20_2 | hypothetical protein |
| TL_JAN20_2 | hypothetical protein |
| TL_JAN20_2 | hypothetical protein |
| TL_JAN20_2 | hypothetical protein |
| TL_JAN20_2 | hypothetical protein |
| TL_JAN20_2 | hypothetical protein |
| TL_JAN20_2 | hypothetical protein |
| TL_JAN20_2 | hypothetical protein |
| TL_JAN20_2 | hypothetical protein |
| TL_JAN20_2 | hypothetical protein |
| TL_JAN20_2 | hypothetical protein |
| TL_JAN20_2 | hypothetical protein |
| TL_JAN20_2 | hypothetical protein |
| TL_JAN20_2 | processive 1,2-diacylglycerol beta-glucosyltransferase |
| TL_JAN20_2 | hypothetical protein |
| TL_JAN20_2 | hypothetical protein |
| TL_JAN20_2 | hypothetical protein |
| TL_JAN20_2 | 5-methylcytosine-specific restriction endonuclease McrA |
| TL_JAN20_2 | hypothetical protein |
| TL_JAN20_2 | hypothetical protein |
| TL_JAN20_2 | hypothetical protein |
| TL_JAN20_2 | hypothetical protein |
| TL_JAN20_2 | hypothetical protein |
